# Estimating interaction matrices from performance data for diverse systems

**DOI:** 10.1101/2022.03.28.486154

**Authors:** Malyon D. Bimler, Margaret M. Mayfield, Trace E. Martyn, Daniel B. Stouffer

## Abstract

1. Network theory allows us to understand complex systems by evaluating how their constituent elements interact with one another. Such networks are built from matrices which describe the effect of each element on all others. Quantifying the strength of these interactions from empirical data can be difficult, however, because the number of potential interactions increases non-linearly as more elements are included in the system, and not all interactions may be empirically observable when some elements are rare.
2. We present a novel modelling framework which estimates the strength of pairwise interactions in diverse horizontal systems, using measures of species performance in the presence of varying densities of their potential interaction partners.
3. Our method allows us to directly estimate pairwise effects when they are statistically identifiable and approximate pairwise effects when they would otherwise be statistically unidentifiable. The resulting interaction matrices can include positive and negative effects, the effect of a species on itself, and are non-symmetrical.
4. The advantages of these features are illustrated with a case study on an annual wildflower community of 22 focal and 52 neighbouring species, and a discussion of potential applications of this framework extending well beyond plant community ecology.

## 1 Introduction

In many biological systems, interactions between system elements (be these species, individuals, etc.) affect population-level performance and together determine the dynamics of the whole system. In order to understand system dynamics when multiple system elements are involved, such complex systems can be represented as networks where the elements are nodes and linked by interactions (Pimm and Lawton 1978). These nodes can take on a wide array of identities, including cells, individuals, populations or species. Likewise, interactions or links can operate via many different mechanisms and have a wide range of effects on the nodes. Biological networks have been observed to typically differ from randomly-assembled networks in important and varied ways, thus informing us on the biological processes structuring these systems (Dunne, Williams, et al. 2002; Kinlock 2019).

Network theory has been widely applied to investigate the structure of biological systems such as food webs and other types of multi-level ecological interaction networks. Because network theory can be used to characterise diversity, stability and other community-level properties emerging from species interactions, it has had a long and meaningful impact on our understanding of ecological communities. There is now a rich body of work describing the structural properties of food webs, plant-pollinator networks, and host-parasite interactions (e.g. Lafferty et al. 2008; Thompson et al. 2012; Dunne, Lafferty, et al. 2013; Stouffer et al. 2014; Cirtwill and Stouffer 2015). Horizontal networks, however, where interactions occur within the same level of organisation (for example interactions between plants belonging to the same food web) have been more neglected by network ecology (Ellison 2019). These species and their interactions have been found to alter the structure, stability and diversity-functioning relationships of the whole multi-level system (Hammill et al. 2015; Giling et al. 2019; Zhao et al. 2019; Miele et al. 2019), which makes integrating horizontal and vertical networks a key challenge in improving our understanding of system dynamics and persistence (Godoy, Bartomeus, et al. 2018).

In many horizontal systems, interactions between species are not always easy to observe empirically and must instead be deduced through other means. A common approach in ecology is to directly quantify the effects of interacting species on the species of interest by evaluating its performance in the absence and presence of potential interaction partners (Connell 1961; Grace and Tilman 1990). ‘Performance’ can refer to any variable of interest that affects the dynamics of the system, for example quantity of resources gathered, biomass, or population growth rate. The resulting interactions are phenomenological and thus not dependent on any specific mechanism, allowing us to capture a wide range of biological processes affecting the dynamics of the whole system (Novak and Wootton 2010). Such methods can quickly become data intensive and computationally complex, however, as the number of species *S* increases and the number of potential direct interactions subsequently increases to *S*^2^. Highly diverse systems pose a further challenge: the abundance distribution of different species is typically skewed, with a few species making up the majority of abundances and a large number of elements remaining rare (Fisher et al. 1943). Given that data collection is limited in time and scope, interactions with rarer species especially may not be observed simply by chance. We thus run the risk of excluding them from analyses regardless of the role they might play (Olesen et al. 2011). Empirically quantifying interaction matrices for diverse horizontal systems thus requires a method that is flexible to both a high number of species, and potential gaps in our records of interactions.

We present a general framework to estimate interactions in diverse horizontal systems. We specify a joint model which allows us to estimate both pairwise identifiable and pairwise unidentifiable interactions from measures of performance in the absence and presence of different interaction partners. We implement the model in STAN, a Bayesian statistical language, and apply it to an ecological case study of an annual wildflower community in Western Australia. Using this dataset, we estimate positive and negative interactions between 22 focal species and 52 neighbouring species and illustrate how to couch these results into established models of population dynamics. This step further widens the potential applications of this novel framework by accounting for other demographic processes affecting species or element performance, and thus system dynamics. We showcase the advantages of this approach by illustrating a range of findings from our case study. We suggest potential applications in species management and conservation that make use of the rich information provided by horizontal interaction networks as developed using our novel framework.

## 2 Methods

We developed a joint modeling framework to estimate pairwise interactions which benefits from several distinguishing features including the ability to estimate both *identifiable* interactions (direct estimates from the observed data) and *unidentifiable* interactions (when observations are missing or too few). After describing the required data (2.1 below), we show how one can estimate identifiable interactions with a unique interaction parameter as described in the neighbour-density dependent model (2.2). We then show how to define and select which interactions are identifiable and which are not (2.3) based on the data available. Independently from the first model, we also describe a response–impact model (2.4) where species have a singular effect on neighbours, and a singular response. This allows us to estimate unidentifiable interactions as the product of element-specific response and impact parameters. Both models contribute to the overall joint model likelihood as detailed in 2.5. Taken together, identifiable and unidentifiable interaction estimates are then used to populate community interaction matrices for the system that describes the effects of all interaction partners on the performance of all focal species.

### 2.1 Data requirements

The joint model framework was initially developed for an ecological dataset where interacting elements (species) affect each other’s performance (lifetime reproductive success). Though we refer to system elements as species throughout this paper, this framework can be applied to data from any interacting group of elements (e.g. cells, individuals, populations, species) which meet the following criteria. First, observations must include some proxy for performance, such as growth (e.g. biomass), fecundity (e.g. number of eggs laid) or chemical production (e.g. oxygen). Second, these observations must also record the identity and density of elements which potentially interact with each focal element. Lastly, observations should be replicated for each focal element with the aim to capture variation in the identities and densities of interaction partners. In addition to these requirements, the experimental design or data may also benefit from observations of focal elements with no interaction partners to better estimate intrinsic performance.

We define *s* as the number of focal species *i*, *i* ∈ {1, .., *s*}, and *t* as the number of interacting species *j* across all *s* focals, *j* ∈ {1, …, *t*}. Typically not all species in the system will be represented in the set of focals, such that *s* ≤ *t*. Measurements of the performance of individual units from each focal element (e.g. seed production of individual plants belonging to a set of focal species) are stored in a vector *p* of length *n* and indexed by *k*, with *k* ∈ {1, …, *n*}. The densities of interaction partners are stored in a matrix *X*, of size *n* × *t*. When an element *j* was absent for a given observation *p_k_*, then *X_k,j_* = 0. Finally, the species identity of each of the *n* focal individuals is stored in the vector *d*, of length *n*, and containing the index of the corresponding species: *d_k_* ∈ {1, …, *s*}.

### 2.2 Neighbour-density dependent model

We quantify the strengths of interactions by regressing the performance of a species - alternatively, a population or any chosen set of replicated units - against the density and identity of other interacting species in a neighbour-density dependent model (NDDM). Increases or decreases in a species’ performance are thus attributed to the changing densities of its interaction partners.

We implement a NDDM for each focal species *i* which regresses the densities of species *j* (*j* = 1, …, *t*) against the measured proxy for performance *p_k_* through a link function *f*(*p_k_*):

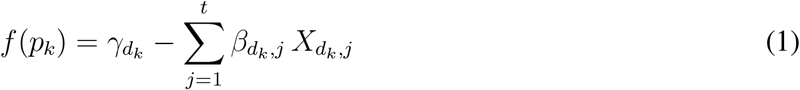

The parameters *β_d_k_,j_* capture the effect of each species *j* on *i* whereas the intercept *γ_d_k__* represents intrinsic performance (in the link scale), a species’ performance in the absence of interactions or when all interaction effects are 0. Note that interacting species *j* can include members of focal species *i* itself, in which case intraspecific interactions are captured by the parameter *β_d_k_,i_*. Furthermore, the equation above places no restrictions on the sign of *β_d_k_,j_*: interactions can be harmful to the focal species (competitive) or beneficial (facilitative). The negative sign in front of *β* does, however, mean that positive interaction estimates should be interpreted as competitive and negative estimates as facilitative. Note that because *d_k_* is the focal species index for each observation, parameters making use of the *d_k_* subscript can use an *i* subscript instead when they are no longer linked to a specific observation of performance.

We implement this model, as well as the RIM described below (in 2.4), as generalised linear models. This means that the relationship between *p_k_* and the right-hand side of the equation does not need to be linear. This can easily be changed by choosing an appropriate link function *f*(*p_k_*) for the data in question. In our case study, for example, we use the link function *f*(*p_k_*) = ln(*p_k_*) since we model our response variable as a negative-binomial variate.

### 2.3 Defining identifiable and unidentifiable interaction parameters

A common issue in observational datasets is that some species or elements may not be observed to interact with many other species or to interact at sufficiently variable densities because data sampling is limited in space and time. This can create a situation in which we cannot estimate all potential interaction parameters (*β_i,j_*) in the NDDM specified above, especially for rare species *j*. For an interaction parameter to be *identifiable*, and thus inferrable by the NDDM, the data must contain measurements of the performance of *i* when interacting with *j* at varying densities. Moreover, the vector of densities of *j* associated with measurements of focal *i* must be linearly independent to all other vectors of densities of other species interacting with *i*. For example, if two species *a* and *b* interact with focal *i* yet have the same density at every measurement or equally proportional densities at every measurement, neither *β_i,a_* nor *β_i,b_* will be inferrable by the NDDM. We define *identifiable* interactions as those which are inferrable following the above assumptions, and *unidentifiable* interactions as those which are not. We construct a matrix *Q* of size *s* × *t*, with *Q_i,j_* = 1 if the corresponding *β_i,j_* parameter is identifiable, and *Q_i,j_* = 0 if not. How we verify linear independance between vectors of neighbour densities and mathematically construct this *Q* matrix is shown in the code shared with the github repository associated to this paper.

### 2.4 Unidentifiable parameters and the response–impact model

Estimating unidentifiable interaction parameters is a complex challenge for which no single answer exists. Here, we address the issue by using an alternative model which assumes that a species *i* will typically have a singular impact (*e_i_*) on and a singular response (*r_i_*) to neighbours independent of neighbour identity (response and impact model, or RIM - see Godoy, Kraft, et al. 2014). A pairwise interaction parameter is therefore the product of a focal species’ *i* response parameter and an interacting species *j* impact parameters. The density-dependent model of performance then becomes:

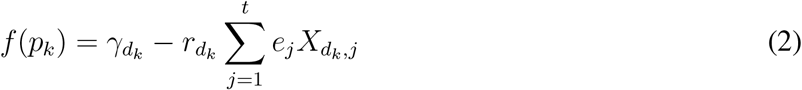

To fit the RIM, the densities of species *j* must be linearly independent to the densities of other species and/or combinations of species, but across the entire *X* matrix. This is in contrast to 2.2 above, where the densities of *j* must be linearly independent within the subset of observations for each focal *i*. Importantly, this is a less strict condition for parameter identifiability. As a result, it is possible to estimate pairwise interactions that would be unidentifiable given Eq. 1 by recognizing that pairwise density dependence in Eq. 2 is given by *β_i,j_* = *r_i_e_j_*.

### 2.5 Joining the two models

The interaction parameters returned by Eqs. 1 and 2 can be used to construct a community interaction matrix *B* of size *s* × *t*, where the effect of species *j* on focal species *i* corresponds to the *i*’th row and the *j*’th column of *B*. An approximation of the matrix of interactions *B* is thus given by:

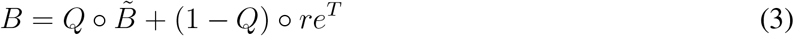

where *Q* is the matrix of identifiable interactions, and 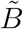 is an *s* × *t* matrix that only has free parameters inferred in the locations where *Q_i,j_* = 1. ∘ represents the elementwise (Hadamard) product, whereby each element *i*, *j* of the first matrix is multiplied by the *i*, *j* element of the second matrix, resulting in a matrix of the same dimension as the operands. In other words, the final interaction matrix *B* takes a free parameter from the NDDM when that parameter is identifiable, and the corresponding RIM estimate, *r_i_e_j_*, when that parameter is not.

We can fit all parameters simultaneously by requiring that both the NDDM and RIM contribute to an overall joint model log likelihood:

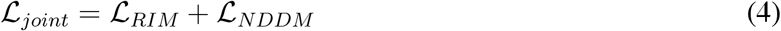

with

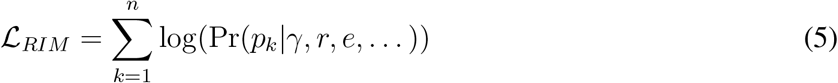

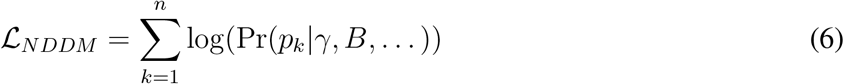

where both sums are across all *n* observations, Pr is the probability density for the model (e.g. negative binomial), *p_k_* the vector of measured performances, *γ, r, e*, and *B* are parameters as defined in Eqs. 1 and 2, and … includes any other parameters (e.g. dispersion if using a negative binomial distribution for the data). Note that *p_k_* and *γ* are common to both the RIM and NDDM.

For a hypothetical dataset with which all interactions are identifiable, then 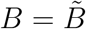 and fitting the RIM is not strictly necessary. The parameter vectors *r* and *e* would still be estimated, but they are independent of the parameters in 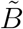 and hence maximization of both likelihoods is independent. Conversely, if no interactions are identifiable then the joint model devolves to the RIM only. In the middle ground—where the majority of datasets are likely to lie—the joint model framework allows us to estimate free interaction parameters when possible, and use *r_i_e_j_* estimates when not. An important distinction to make is that the NDDM estimates identifiable interactions only, whereas the RIM estimates all interactions, pairwise identifiable and pairwise unidentifiable. However, by maximising Eq. 4 we allow both models to provide good fits to the data but also for *r* and *e* to “adjust” around inferred values of 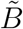.

### 2.6 Making interaction parameters comparable across focal species

The *β_i,j_* estimates returned by the framework describe the effect of species *j* on the performance of species *i*. Differences in the magnitude of these interaction terms may thus reflect intrinsic differences in performance, which can vary in time, space and between species. Two species may for example have different baseline values of reproductive fitness. In order to make the effects of interactions comparable between species, the interaction terms returned by both models above can be transformed into scaled interaction strengths (Laska and Wootton 1998). The appropriate scaling is determined by rewriting the neighbour density-dependent model (Eq. 1) into a form equivalent to a Lotka-Volterra competition model:

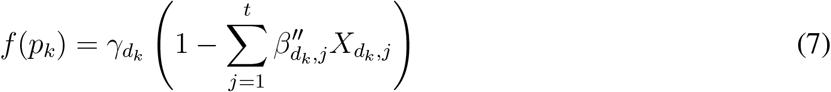

This reveals that our interaction terms can be rescaled into interaction strengths by dividing them by the recipient element’s intrinsic performance:

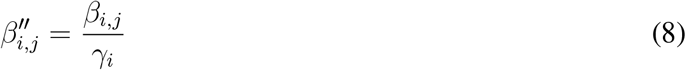

Though this scaling step is not strictly necessary, for ecological datasets these scaled interaction strengths have the benefit of being directly comparable both across species and across environmental contexts where reproductive fitness may vary (Wootton and Emmerson 2005), leading to a wider range of potential applications.

### 2.7 Integrating interaction strengths into models of population dynamics

In certain instances, models of species population dynamics can be used to further extend the usefulness of the framework presented here. We suggest two cases where such an application may be useful. Firtly, the variable chosen to measure the performance of focal species *p* may not directly translate into a measure of performance which is relevant to system dynamics, due to inherent practical constraints with collecting empirical data. For example, the life-history reproductive strategies of certain species may lead to measures of high fecundity (or performance) in the field which do not account for low survival rates post-observation (Broekman et al. 2020). In these cases, population dynamics models can be used to account for species-specific demographic rates into estimates of interaction effects. Alternatively, we might be more interested in the effects of interacting species on the density or growth rate of a focal species rather than on it’s performance. In this scenario, a population dynamic model can be used to translate interaction effects on the measured variable into interaction strengths affecting the variable of interest.

In both cases, an established population dynamic model is required as well as knowledge of any crucial species-specific demographic rates. This step is illustrated for our case study in the Supplementary Methods 6.2.3, where an annual plant population dynamic model is used to transform effects on wildflower seed production (the measured proxy for performance) into effects on population growth, and includes species-specific estimates of seed germination and survival rates.

### 2.8 Model fitting

We implement the NDDM (Eq. 1), the RIM (Eq. 2), and the joint model as generalised linear models in STAN (Carpenter et al. 2017), a Bayesian statistical language where coefficient values will be estimated by MCMC sampling. Using STAN requires translating the model formula into the STAN language, setting priors for parameters to be estimated, and using an indexing system to discriminate between identifiable and unidentifiable interactions. We provide a working example of the STAN code used to specify and set this model to the case study below, as well as the necessary R scripts and functions to run the model on a simulated dataset on our public github repository https://github.com/malbion/JointModelFramework. From the model file, only the link function for *p_k_* and it’s parameterisation need to be modified in order to apply it to a differently-distributed dataset. Additionally, non-integer measures of performance (e.g. biomass) should be redefined as real rather than integers in the data block. In the code given, a negative binomial distribution is used to fit seed production but a different distribution may be more appropriate when using other measures of performance.

Bayesian models are often run on multiple MCMC chains. In our case, we deliberately run only one MCMC chain because the latent variables in our model (*r_i_* and *e_j_*) are invariant to sign switching. This means that different MCMC chains can return coefficient values which are of the same magnitude, but opposing signs. As with the boral function in the boral R package (Hui 2021) and the MCMCfactanal function in the MCMCpack package (Martin et al. 2011), we run one chain only to simplify issues with checking chain convergence and computing parameter estimates which arise from sign-switching. Though many MCMC convergence diagnostics require multiple chains to be computed, single chain convergence can be assessed with the Geweke convergence statistic (Geweke 1992), for example with the geweke.diag() function from the coda package (Plummer et al. 2006), as well as visually checking traceplots.

STAN returns parameters as distributions which maximise the likelihood, and are conditioned by the data and priors. Priors describe the distribution of plausible values which these parameters may take. For an introduction to Bayesian inference which relates the use of priors to frequentist hypothesis testing, see Ellison (1996). We recommend investigators experiment with setting different informed priors to both improve model convergence and verify the robustness of parameter estimates. The resulting parameters are termed posterior distributions, and samples from the posterior are drawn for analysis. Using parameter distributions rather than point estimates allows for easy inclusion of uncertainty in the analysis of results, we therefore recommend bootstrap sampling from each posterior interaction strength distribution to create multiple samples of the community interaction matrix. Ellison (2004) also provides an accessible review of parameter estimates and the use of posterior distributions using a worked example on ant species richness data.

### 2.9 Case study

We applied this framework to an annual wildflower community dataset from Western Australia collected in 2016. This dataset contains over 5000 observations of individual plant seed production from 22 different focal species, and the identity and density of all neighbouring individuals within a 3 to 5 cm radius of the focal individual. We used the framework to quantify interactions between these 22 focal species and 52 neighbour species, and derived scaled interaction strengths with a well-supported population dynamics model for annual plants with a seed bank (Levine and HilleRisLambers 2009; Bimler et al. 2018) which required experimentally-measured species demographic rates. Viable seed production was used as a measure of performance and modeled with a negative binomial distribution and a log link. Further details on the data, the model fitting and the procedure for deriving scaled interactions from the population dynamics model are available in the Supplementary Methods 6.2.

## 3 Results

The joint model framework returns a matrix *B* which quantifies the effect of interacting species *j* (columns) on the performance of focal species *i* (rows). The interactions (values) which make up *B* can be positive or negative, non-symmetrical (the effect of element *i* on *j* does not necessarily match the effect of element *j* on *i*) and include intraspecific effects (the effect of element *i* on itself). We illustrate the advantages of this approach in the case study results below.

### 3.1 Case study results

The model returned estimates for all 1144 interactions between 22 focal species and 52 interacting species, of which 56.7% were identifiable and estimated by the NDDM. When accounting for interactions between focal species only, 82.0% of interactions were identifiable. We conducted a posterior predictive check comparing simulated performance data from the joint model to observed values (Figure 1), this is especially important for verifying that the appropriate distribution and link function are being used for the data at hand. The joint model also returns simulated performance data for the RIM only, which we also checked visually (Supplementary Results 7 Figure 1). Model parameters were sampled 1000 times from the 80% posterior confidence intervals returned by STAN to construct our parameter estimates. Interaction estimates were scaled according to focal intrinsic fitness into interaction strengths comparable between focal species, and we applied bootstrap sampling from each resulting interaction strength distribution to create 1000 samples of the scaled community interaction matrix.

**Figure 1:**
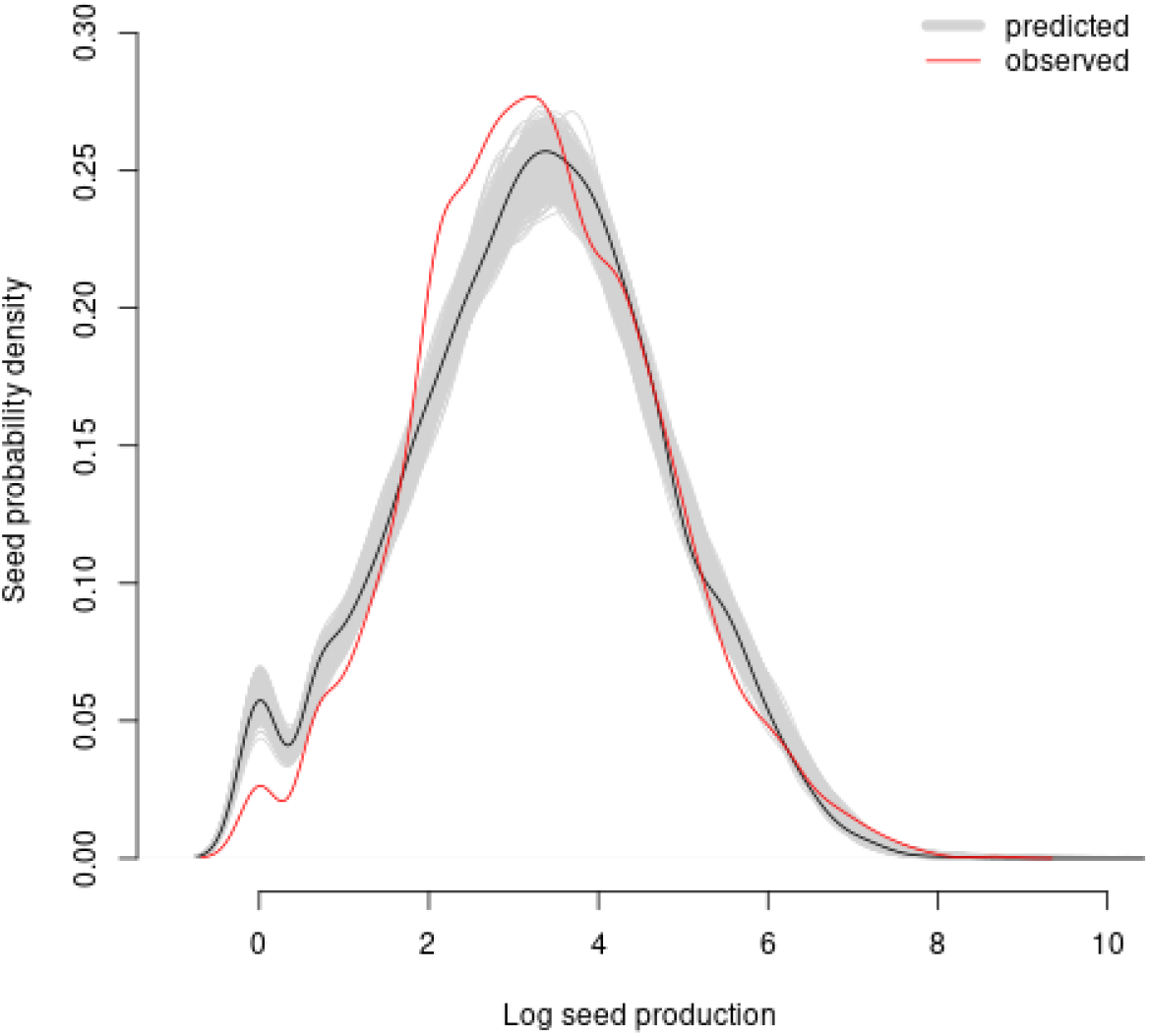
Posterior predictive check showing the density distribution of observed seed production values (red line) to simulated seed production values (light grey) as estimated by the joint model, on a log scale. Simulated values were generated using the 80% posterior confidence intervals for each parameter, the black line shows simulated values using the median of each parameter.

For our case study, the resulting community interaction matrix was non-symmetrical and included both positive (competitive) and negative (facilitative) values, as shown in Figures 2.A & B. Here we represent the community matrix as a network between all 22 focal species, taking the median value of each scaled interaction across all samples. Overall, 63.6% of focal species had a median interspecific interaction strength which was competitive, making competition the dominant interaction type. The median value of 46.7% of focal × neighbour interactions, however, were facilitative, as were the medians of 43.3% of all focal × focal interactions. As a result, 47.6% of interactions between pairs of focal species were of opposing signs such that *i* competes with *j* but *j* facilitates *i*. Furthermore, the elements of the diagonal (the effect of a species on itself) were able to be estimated, which allows us to quantify how much a species regulates it’s own performance. For 11 of our 22 focal species, the scaled distributions of these intraspecific effects did not overlap with 0, which suggests individuals of those species have a non-trivial effect on other individuals of the same species.

**Figure 2:**
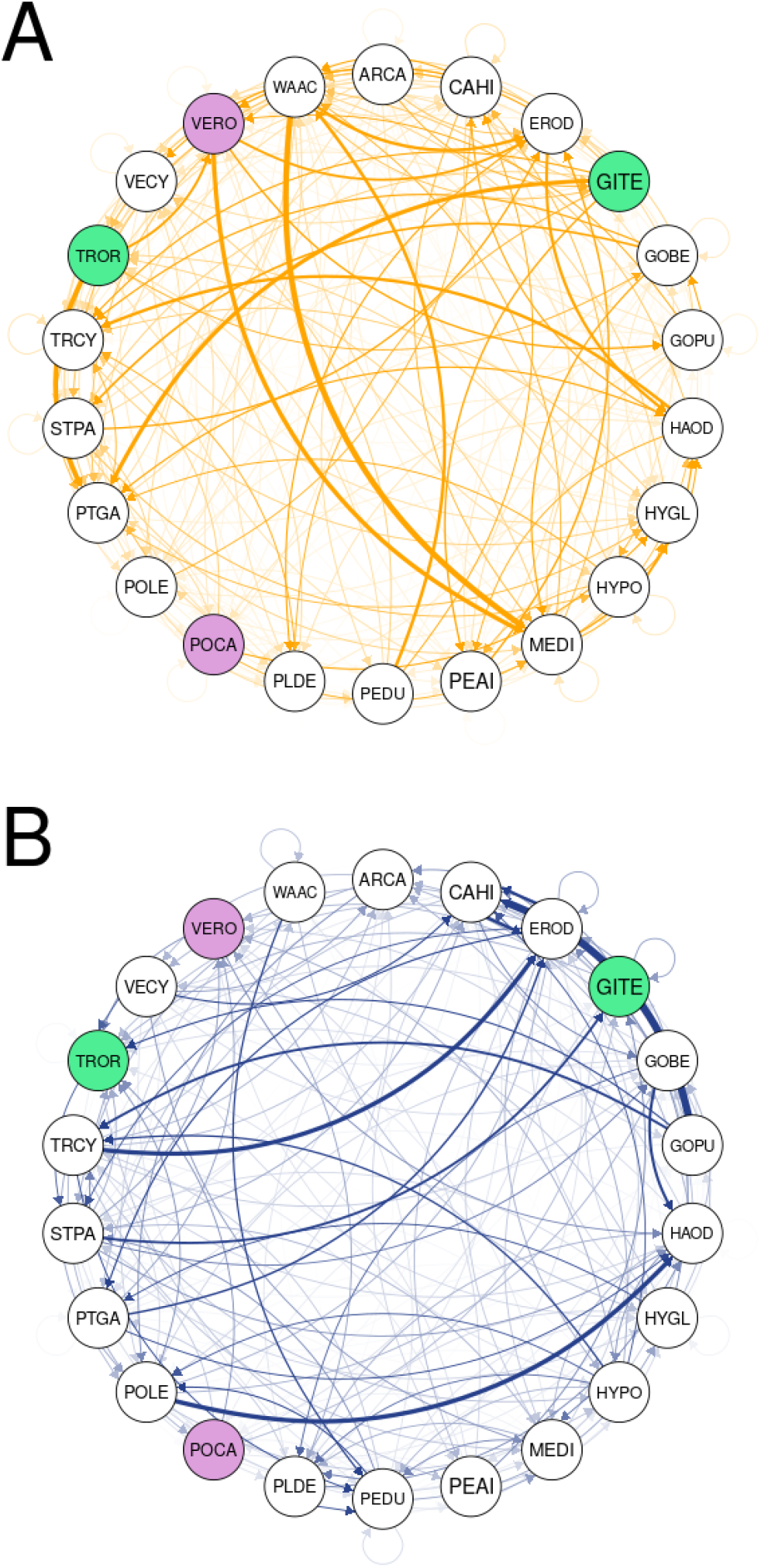
Competitive (A) and facilitative (B) scaled interaction networks estimated from our model framework. Competitive and facilitative interactions (A and B) are here shown separately for ease of view but were analysed together. Only focal species are included in these networks, arrows point to species *i* and line thickness denotes interaction strength. Interaction strengths are given as the median over 1000 samples. Purple coloured nodes correspond to highly abundant native species, whereas green nodes indicate potential keystone species, as further described in Figure 3.

### 3.2 Examples of ecological applications

We illustrate a few potential applications of our framework by exploring questions of common ecological relevance and how these can be answered using the results from our case study. Each question below highlights some of the advantages of our resulting interaction network: intraspecific interactions, non-symmetrical interactions, and the ability to estimate positive and negative interactions. We were unfortunately unable to compare our results to other methods for estimating such a diverse set of plant-plant interactions because there are currently none other that we are aware of that do this.

#### Do abundant natives under-regulate their population density compared to rarer native species?

One hypothesis as to why certain plant species are more abundant than others is that they tend to compete with themselves less strongly than rare species (Yenni et al. 2012; Yenni et al. 2017). Hypothetically, this release from intraspecific competition pressure allows them to reach much higher densities than species which strongly compete with themselves. In our case study, we can explore this hypothesis by plotting the effect of a species on itself (the diagonal of the community interaction matrix, or intraspecific interactions) against it’s density as in Figure 3.A. Intraspecific interactions are at their weakest when close to 0. The two most abundant native species *Velleia rosea* (VERO) and *Podolepsis canescens* (POCA) highlighted in purple fall very close to the median intraspecific interaction strength. This suggests that *V. rosea* and *P. canescens* do not reach high densities through an under-regulation of their population density but through other means (e.g. access to a larger niche space).

**Figure 3:**
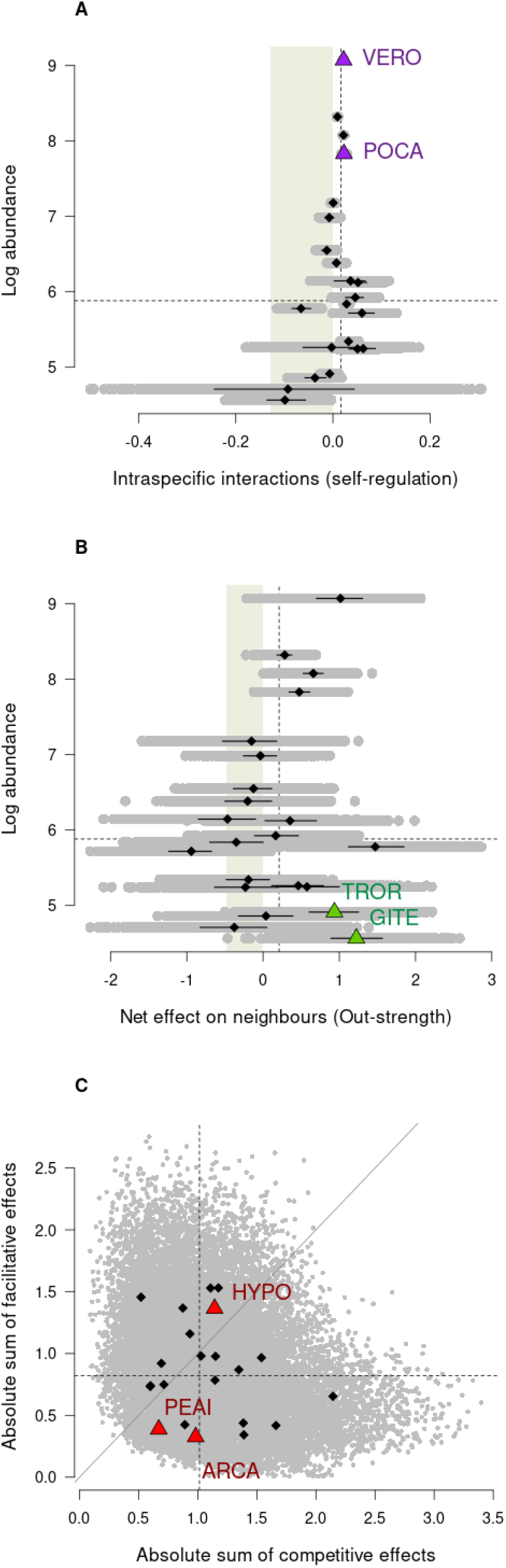
Understanding interaction effects can help identify species of particular ecological importance to a system. For all graphs, diamonds are species medians across all network samples, black lines cover the 50% quantile and grey dots indicate the full range of out-strength values as calculated from 1000 sampled networks. The shaded part of the graphs show facilitative (negative) interactions. Dashed lines represent the median value for all focals. Coloured triangles indicate the species refered to in the main text for each of the ecological questions associated with (A), (B) and (C). In (A), the x-axis shows the strength of scaled intraspecific interactions, that is how strongly a focal species interacts with itself, plotted against a focal species’ total log abundance (y-axis). Values over 0 indicate competition, and values less than 0 indicate facilitation. The two most abundant natives, *Velleia rosea* (VERO) and *Podolepsis canescens* (POCA) in purple, do not seem to compete with themselves any more or less strongly than the median for all species in the system (dashed line). (B) shows the sum of interaction effects of focal species on neighbours (x-axis) against the focal species’ total log abundance (x-axis). On the x-axis, values greater than 0 indicate that a focal species has an overall competitive effect on neighbours, and values less than 0 indicate that it has an overall facilitative effect. Green diamonds identify species with low overall abundance but strong competitive effects on neighbours: *Trachymene ornata* (TROR) and *Gilberta tenuifolia* (GITE). (C) decomposes a focal species’ net interaction strength into its competitive effects (x-axis) and facilitative effects (y-axis). Red diamonds show the exotic species *Hypochaeris glabra* (HYPO), *Arctotheca calendula* (ARCA) and *Pentameris aroides* (PEAI). The light grey diagonal shows where x = y, species above that line have an overall facilitative effect on other species whereas those below that line have an overall competitive effect.

#### Which species may be taking on keystone roles in the system?

Keystone species have strong effects on the dynamics of the whole ecosystem, such that their exclusion from a community can create significant changes in species density and composition (Paine 1969). Furthermore, the impact of keystone species on other species is disproportionately large relative to their density (Power et al. 1996; Piraino et al. 2002; Libralato et al. 2006). The keystone species concept is relevant to ecosystem management and conservation in helping identify species of particular importance for the safeguarding of a whole system (Soule et al. 2005). Though determining which species truly serve keystone roles has historically involved extensive ecological experimentation (e.g. Paine 1992), we can identify potential candidates by comparing a species’ impact on the population growth of other species to it’s own density (Libralato et al. 2006). It is important to note that because our framework allows for asymmetrical interactions, we are able to differentiate a species’ impact on other species from it’s response or sensitivity to neighbours (Broekman et al. 2020). Figure 3.B highlights two native species in green which may be potential keystone species due to having strongly competitive effects on the rest of the community overall, despite low densities: *Trachymene ornata* (TROR) and *Gilberta tenuifolia* (GITE).

#### Do all exotic species compete with native species?

Though many invasive species compete with natives (Naeem et al. 2000; Corbin and D’Antonio 2004; Riley et al. 2008; Zheng et al. 2015), several studies have found evidence of invasives facilitating natives, with cascading effects on other species and net positive effects on ecosystem processes (Rodriguez 2006; Ramus et al. 2017; Wainwright et al. 2019). By allowing for positive and negative interaction strengths between species in a system, we can determine which exotics are harmful or beneficial to a native’s performance. Figure 3.C plots the sum of a species’ competitive effects on neighbours against the sum of it’s facilitative effects on neighbours. Exotic species are identified in red. Out of these three exotic species, two have overall competitive effects on the community: *Arctotheca calendula* (ARCA) and *Pentameris aroides* (PEAI). *P. aroides* interacts weakly with neighbours, whether it is competing or facilitative, and *A. calendula* does compete but still lies below the median sum of competitive effects for the community, it’s overall competitive effect being largely driven by weak facilitation. *Hypochaeris glabra* (HYPO) on the other hand has strong effects on other species, both competitive and facilitative, and has an overall facilitative contribution to the community. The results from our case study suggest that here at least, the effects of exotic species on native species are complex and species-dependent.

## 4 Discussion

Our novel framework quantifies the effects of interacting species and reciprocal performance, allowing the estimation of diverse, horizontal interaction matrices. The resulting matrices are non-symmetrical and can contain both positive and negative interactions, as well as the effect of a species on itself. This framework is flexible to metrics of performance, group identity (e.g. species, population, etc.) and diversity. We also propose a way to estimate unidentifiable interactions from those which are identifiable. The matrices generated through this framework can be transformed into interaction networks through the use of further models describing the system’s interaction dynamics. These features make it particularly useful in an ecological context, as illustrated in our case study on a diverse wildflower community, as well as flexible for use with data from the wide range of complex systems dominated by horizontal interactions.

Here, we illustrate the unique useful features of this framework using a typical plant-dominated ecological system for context. In our case study, we show how wildflower species are linked through plant-plant interaction networks. In turn, this network can help us identify what roles specific species play within a community, and explore how the mechanisms maintaining diversity and stability operate in these systems. By estimating the effects of species on each other’s performance, and subsequently their population growth and patterns of density, our method stands in contrast to association networks and returns qualitatively different information. There are a wide range of association network frameworks used in ecology (Burns and Zotz 2010; Losapio, Cruz, et al. 2018; Montesinos-Navarro et al. 2018) and collectively, they are a common alternative to estimating interaction networks in high-diversity systems as they capture spatial associations between species (e.g. Saiz et al. 2018) and require easier-to-obtain data. Association networks also benefit from easy implementation with a wide range of packages in R (e.g. Griffith et al. 2016) though the resulting networks are typically symmetrical and cannot capture a species’ effect on itself. Moreover, association networks remain poor predictors of species interactions and rarely match empirical estimates (Sander et al. 2017; Barner et al. 2018; Thurman et al. 2019; Blanchet et al. 2020).

Species interaction networks have a wide range of practical applications, such as evaluating ecosystem response to human-altered landscapes, guiding future management decisions (Ross et al. 2011) or exploring how communities may respond to global warming (Gorman et al. 2019). Conservation and ecosystem management efforts aimed at regulating species abundances can, for example, use the information provided by an interaction network to prioritise which species to conserve or eradicate based on their role in the community. Such roles can be deduced by a species’ position in the interaction network (Cirtwill, Dalla Riva, et al. 2018) and as illustrated in our case study. Identifying keystone, foundation and other important types of species roles is also helpful for understanding biological diversity, ecosystem integrity and functioning, especially in response to disturbances and other stresses (Nyakatya and McGeoch 2008; Orwin et al. 2016; Losapio and Schöb 2017; Narwani et al. 2019). The examples we describe in our case study are not exhaustive, but serve to illustrate how interaction networks can help us understand both community dynamics overall and the effects & response of specific species towards the community.

Quantifying the matrix of scaled interaction strengths between species in horizontal communities can also allow us to explore how the mechanisms maintaining diversity and stability operate in these systems and across a broad number of species. Self-regulation, for example, is an extremely important driver of community stability (Barabás et al. 2017) and arises from how individuals of the same species interact with one another. Whereas same-species effects cannot be estimated by association networks, our framework quantifies intraspecific interactions and allows us to measure the strength and prevalence of competitive and facilitative density-dependence. Measures of intra and interspecific interactions can also allow us to estimate niche overlap between species (for an example, see Chu and Adler 2015); weak interactions between species suggest that they are not sharing or competing for many resources, and thus may have large niche differences in the community.

Another key feature of our model framework is the inclusion of facilitative interactions, which have traditionally been disregarded in plant population models and theoretical frameworks of plant diversity-maintenance. The importance of facilitative interactions to community structure and patterns of abundance has long been recognised (Callaway et al. 1997) but there is still little consensus on how they may affect biodiversity (Bruno et al. 2003; Brooker et al. 2008). Recent work suggests facilitation may be more widespread than traditionally thought (Gross et al. 2015; Picoche and Barraquand 2020) and can benefit species diversity and stability depending on the circumstances (Coyte et al. 2015; Brooker et al. 2008). Our framework provides a means to investigate the prevalence and strength of facilitation across multiple species, and how it may act in relation to competition and species diversity.

Ultimately, quantifying species interaction networks allows us to apply tools from network theory which will help us understand not only how species interact, but also how these interactions drive community-level patterns of density and diversity. Several metrics already exist for describing horizontal network structure such as weighted connectance (Ulanowicz and Wolff 1991) or relative intransitivity (Laird and Schamp 2006), though these are fewer than for trophic or unweighted networks networks (e.g. Bersier et al. 2002; Delmas et al. 2019). Adapting measures of nestedness or modularity for example to non-sparse networks (as as horizontal communities typically are) would allow us to further characterise how interactions and species are organised. These metrics relate to various aspects of stability and could greatly inform us on how diversity is maintained between species belonging to the same trophic level. Likewise, networks also provide several ways of measuring and describing species roles in their respective communities (Cirtwill, Dalla Riva, et al. 2018) for example through the use of structural motifs, unique patterns of interacting species which together make up the whole network. Motifs have been found to have important biological meaning in food webs (Bascompte and Melian 2005) but remain to be identified for single-trophic and bipartite networks.

### 4.1 Limitations

Our novel framework includes a versatile approach for estimating interactions between species which are not observed to cooccur. Identifiable interactions are estimated with a unique parameter, as opposed to the latent variables used to quantify unidentifiable interactions. Care should therefore be taken to assess the different assumptions behind our estimatation of identifiable and unidentifiable interactions, and how well these may apply to any particular system. It is also important to consider the likely reasons for unidentifiable interactions. Forbidden links are a subset of potential interactions which cannot be observed, often due to physical constraints (e.g. biological mismatch) or spatio-temporal uncoupling. For example, a pair of short-lived annual plants might have such opposing phenologies that their growing seasons never overlap in the field. We direct the reader towards the literature on forbidden interactions (Olesen et al. 2011; Jordano 2016) for solving these cases.

Our model framework is aimed to helping empiricists who would like to estimate species interactions in non-trophic communities. Though the MCMC sampling algorithm does allow for many parameters to be estimated, it is crucial to check chain convergence and model behaviour to verify that the full parameter space is sampled. The amount of data required for sampling diverse communities may still be substantial. Grouping species which appear very rarely is also a common strategy to avoid over-parameterisation. In our case study, neighbour species which were recorded fewer than 10 times across all observations were grouped into an ’other’ category with its own interaction effect. If interactions with or between very rare species are the explicit object of a study, however, data-collection should focus on amassing observations of focal individuals from those rarer species to be able to estimate interaction strengths.

### 4.2 Conclusion

There is now a rich body of work describing the characteristics of food web, plant-pollinator and host-parasite interactions, but fewer network approaches focus on non-trophic interactions such as those occurring between plants (Ellison 2019). Here we present a novel framework which makes the process of inferring interactions in horizontal systems easier, whilst allowing for many and multiple types (competitive and facilitative) of interactions. In turn, this allows the application of network theory tools to the management of non-trophic systems, as well as to deepening our understanding of diversity, stability and other community-level properties which emerge from interactions. By applying our framework to a wildflower case study, we find that over-abundant species do not appear to self-regulate more strongly than rare species, contrary to expectations. We also identify species which have strong effects on their neighbours, and which exotic species may be more of a threat to the native community. Though we illustrate our study with one particular ecological dataset, the method presented here could be adapted for use on a wider array of horizontal systems such as those found in microbial, neural, and social networks.

## Acknowledgements

We thank M. Raymundo & I. Towers for the seed rate data used in this paper, C. Bowler, T. Britton, and J. Ikin for their work in sample processing for this data set, as well as Rob Freckleton and Jacopo Grilli for insightful comments on the manuscript. We would also like to thank Stefano Allesina and two anonymous reviewers for their valuable comments on an earlier version of this manuscript. We also wish to acknowledge the Traditional Custodians of the Country from which the case study data was collected, the Yamatji People, as well as those of the land on which this research was conceived of, carried out, and written, the Jagera and Turrbal Peoples. This work was made possible by funding awarded to M.M.M. (DP170100837) by the Australian Research Council. D.B.S. is grateful for the support of the Marsden Fund Council, from New Zealand Government funding (grant 16-UOC-008).

## Competing interests

The authors declare no competing interests.

## 6 Supplementary Methods

### 6.1 Model code and parameterisation

We make the several observations in addition to comments in the code. Firstly, identifiable interactions (beta ij in the code) are defined as a vector, which must then be matched to their correct position in the interaction matrix. This is the role of the istart, iend, icol, and irow vectors defined in the data block. Our github repository also contains the data _prep.R file, which will show how to calculate these vectors from the input data. Secondly, we impose the following constraints to improve convergence, avoid overparameterisation and maintain identifiability of our parameters *r_d_k__* and *e_j_* (Huber et al. 2004; Kidziński et al. 2021; Niku et al. 2021). We define the effect parameters as a unit vector, which means we only require K-1 degrees of freedom (where K is the total number of neighbour elements) to estimate all effect values. This loss of a degree of freedom arises from the fact that the matrix of *r_d_k__ e_j_* parameters is of rank 1. The first response parameter is also forced to positive. This improves convergence by providing an anchor for all other parameter values to ’rotate’ around. Though these latter two have implications for our estimates of the latent variables *r_d_k__* and *e_j_*, estimates for identifiable and unidentifiable interactions should not be affected.

### 6.2 Case study Methods

#### 6.2.1 Community data

We applied this framework to annual wildflower community dataset from Western Australia. This system is a diverse and well-studied community of annual plants which germinate, grow, set seed and die within approximately 4 months every year. Individual fecundity data were collected in 2016, when 100 50 × 50 cm plots established in the understory of West Perenjori Reserve (29°28’01.3”S 116°12’21.6”E) were monitored over the length of the full field season. The resulting dataset includes between 29 to over 1000 counts of individual plant seed production from 22 different focal species (with a median of 108 observations per species), in addition to the identity and densities of all neighbouring individuals within the interaction neighbourhood of each focal plant. Interaction neighbourhoods varied in radius from 3 to 5 cm depending on the size of the focal species (Martyn 2020). Total neighbourhood diversity was 71 wildflower species, 19 of which were recorded fewer than 10 times across the whole dataset. The species-specific effects of this latter group of species on focals were deemed negligible due to their extremely low density, they were thus grouped into an ’other’ category and their effects on focals averaged. This resulted in 53 potential neighbour identities. Half of all plots were thinned (a quarter to 60% diversity and a quarter to 30%) to mitigate possible confounding effects between plot location and plant density, and thinning did not target any particular species.

In study systems which allow it, a proportion of interaction neighbourhoods can instead be thinned prior to the experiment to randomly remove neighbouring individuals and provide observations for low-density estimates of interactions. Though this steps is not strictly necessary, thinning certain neighbourhoods can also reduce potential confounding effects between the environment and interactions and thus provide more accurate estimates of interaction effects. Environmental data known to affect performance can also be recorded and included in the model (as a random effect for example) to minimise those confounding effects.

We required species demographic rates (seed survival and germination) in order to scale model interaction estimates into interaction strengths. Species demographic rates for 16 of our focal species were estimated from a database of field experiments carried out between 2016 and 2019 where seedbags were placed in the field to estimate germination rates, and ungerminated seeds were evaluated in the lab for survivability. The remaining species were assigned mean demographic rates from these experiments. Further details on the methods used for collecting those seed rates are available in section 6.2.4.

#### 6.2.2 Model fitting

We fit the model using R version 3.6.3, STAN and the rstan package (R Core Team 2020; Carpenter et al. 2017; Stan Development Team 2020). Estimates of seed production were fit with a negative binomial distribution. The model was run with 1 chain of 10000 iterations, discarding the first 5000. Models were checked for convergence using the geweke.diag() function from the coda package (Plummer et al. 2006) and traceplots were visually inspected to verify good chain behaviour. Model parameters were sampled 1000 times from the 80% posterior confidence intervals to construct our parameter estimates. We then applied bootstrap sampling from each resulting interaction strength distribution to create 1000 samples of the community interaction network.

#### 6.2.3 A model for annual plant population dynamics

The above model framework returns species-specific estimates of intrinsic fitness (*γ_d_k__*), as well as as a species x neighbour matrix *B* of identifiable (*β_d_k_,j_*) and unidentifiable (*r_d_k__e_j_*) interaction estimates which quantify the effects of one neighbour *j* on the intrinsic fitness of a focal species *i*. Though useful as they are, these estimates can lead to a wider range of potential applications when integrated into models of population dynamics. For example, we might be more interested in the effects of neighbours on the density or growth rate of a focal species rather than on it’s proxy for lifetime reproductive success. Importantly, it is necessary to specifiy a model describing population dynamics in order to draw conclusions about the effects of interactions and network structure on the maintenance of community diversity and stability.

We defined the following model for annual plants with a seed bank (Levine and HilleRisLambers 2009; Mayfield and Stouffer 2017; Bimler et al. 2018) which describes the rate of change in a focal species’ *i* abundance of seeds in a seed bank from one year to the next:

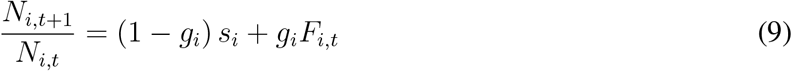

where *F_i,t_* measures the number of viable seeds produced per germinated individual whilst *g_i_* and *s_i_* are the seed germination and seed survival rate, respectively. In a simplified case where the focal species *i* interacts with only one other species *j*, our use of a log link function implies that *F_i,t_* in this model of population dynamics is given by:

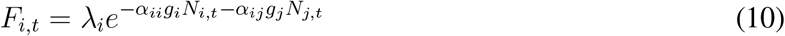

where λ*_i_* corresponds to seed number in the absence of interaction effects, and *α_ii_* and *α_ij_* are the interaction strengths between species *i* and its intraspecific and interspecific neighbours respectively. Here it is *α_ij_* and *α_ii_* which are equivalent to *β_d_k_,j_* in Eq. 1 of the main text. We determine the scaled interaction strengths *α*″’s by including λ*_i_*, *g_i_* and *s_i_* in such a way that these variables are cancelled out when the *α*″’s are substituted for the *α*’s in our annual plant population model (Godoy and Levine 2014; Bimler et al. 2018).

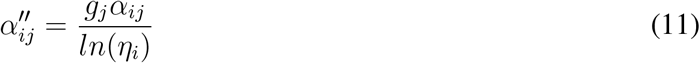

with 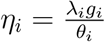 and *θ_i_* = 1 – (1 – *g_i_*)(*s_i_*). Note that our model evaluates the rate of change of seeds in the seed bank, and this is reflected in the scaling terms used to compare interaction strengths between focal species. Substituting *α*″’s for *α*’s in Eq. 9 gives us:

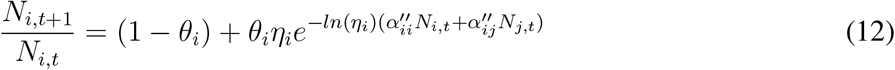

where we can see that the *α*″’s are directly proportional to the density of neighbours. Relating this population model to the joint model framework, we recover the following:

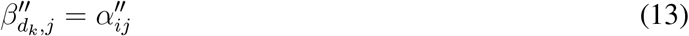

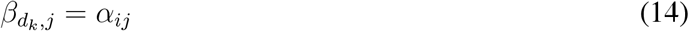

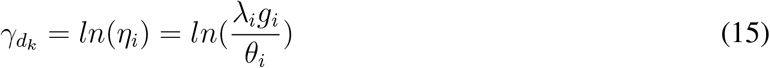

As we show here, the exact form of the rescaled interactions as well as intrinsic fitness can therefore vary depending on the specific population dynamic model applied and may include other demographic rates which reflect species-level differences in growth and mortality. Because intrinsic fitness is estimated by the model framework and not directly observed, we used the mean of the *γ_d_k__* posterior distribution returned by our model in our scaling of the interaction coefficients.

#### 6.2.4 Seed germination and survival data

Seed demographic rates were collected from a set of field experiments conducted by T. Martyn, M. Raymundo and I. Towers at Perenjori reserve between 2015 and 2019. Experiments differed both in the methods and in which focal species were included in ways which are detailed further below, such that each focal species had a different number of replicates across all experiments. Given how much seed rates estimates have been found to vary within species and according to a range of both individual and environmental factors, we chose to average results from these multiple experiments for each focal species in order to provide a point estimate which captures a wide range of conditions under which seeds may grow. For those species which did not have any field estimates of seed rates (*Austrostipa elegantissima*, *Erodium sp*., *Petrorhagia dubia*) or seed survival rate (*Gilberta tenuifolia*), no replication (*Waitzia acuminata)* or an unrealistically low estimate of germination rate (*Goodenia pusilliflora)*, we substituted the community mean instead.

For each experiment, mature seeds were collected at the end of the growing season (September - October) from multiple populations of each focal species located throughout the reserve. Immature or damaged seeds were not included, and collected seed was homogenised for each focal species to elimiate bias associated with local adaptation within populations. Germination rate was estimated by planting seeds in the field along gradients of soil phosphorus, woody canopy cover and herbaceous vegetation density and either directly counting the number of seeds which had germinated or comparing recruitment rates to unplanted plots after a sufficient amount of time had elapsed. Seeds were planted during late-September to mid-October, mimicking natural seed dispersal timing for wildflowers in the area. Seed surival rates were estimated using either the remaining seeds or a separate batch of seeds and assessing viability of the seeds using tetrazolium staining.

##### T. Martyn experiment

For 19 focal species (the full species list excluding *A. elegantissima, P. dubia* and *G. tenuifolia)*, germination bags containing 20 seeds each were planted in the field in 2016 across multiple areas of Perenjori Reserve. Out of the 30 bags, 19 were collected in 2017 and the remaining were collected in 2018. Due to a severe drougth in 2017, half of the bags collected that year were watered during the field season and prior to collection. Germination bags were then brought back to the Mayfield Lab facilities at the University of Queensland, Brisbane, and seeds extracted. Seeds were examined for signs of germination in the field (broken or empty seed coat) and those remaining were placed in germination trays and a germination chamber to mimic light and temperature conditions conducive to germination. Trays were watered with Gibberellic acid once to twice a week and seedlings were recorded and removed until no more seedlings emerged. Remaining, ungerminated seeds were then assessed as dead (moldy) or potentially viable. The remaining potentially viable seeds were assessed for viability using tetrazolium staining, contributing to our estimates of seed survival rates. For this procedure, embryos in each seed were exposed by either removing the seed coat or by creating a thin cut along the seed coat. The exposed embryos were then placed on a six-well germination plate and 2 ml of 0.25% Tetrazolium solution was added to each well to stain the embryos, before covering them and storing them at 25°C overnight. To check for staining, embryos were dissected under a dissecting microscope. Viable seeds showed a dark pink embryo while non-viable seeds did not stain or were stained in a splotchy way.

##### M. Raymundo experiment

This experiment was carried out on the focal species *H. glutinosum*, *T. cyanopetala*, *T. ornata* and *V. rosea* from 2015 to 2017. However, a severe drought in 2017 made the second round of data collection impossible and thus we only include results for 2016 here. Ten plots were established measuring 0.5 m x 0.5 m at each of three sites in Perenjori Reserve for a total of 30 plots. Each plot was divided into 25 0.1 m x 0.1 m subplots and focal species were randomly assigned a subplot in each plot. Thirty seeds of each focal species were planted in the designated subplot in late September 2015 and a plastic ring 10 cm in diameter and 1 cm high was placed in each subplot where seeds were added to limit seed movement among subplots. Another five subplots were assigned plastic rings to serve as controls for the effect of the rings on non-experimental communities. The remaining 15 subplots served as controls where no seeds or rings were added allowing for recruitment from either natural dispersal or from the seed bank. Blocks were placed in such a way as to span shaded and open areas, bare ground and dense herbaceous vegetation, and areas with native dominated and exotic dominated assemblages. Before implementing the experiment in 2015, plots were surveyed to record the number and identity of all adult plants in each subplot. Due to the randomization of seed addition into subplots, some subplots had focal species already in them. As all focal species were common to this reserve, it is also likely that seeds for all species were in the seed banks in at least some subplots. There was no way to determine this in advance, though when adult individuals of a focal species were present in a subplot prior to the implementation of our experiment, we expected that some seedlings in the following year would be from the seedbank as well as our planted seeds and looked for evidence of this (more than 30 individuals) in data from 2016. We therefore compared average densities of successful focal recruits and those which emerged in situ between sown, control, and ringed subplots to assess seed limitation and germination rate. To measure seed survival rates, thirty seeds of each focal species were also assessed for viability using tetrazolium staining using the same procedure as for the T. Martyn experiment.

##### I. Towers experiment

This experiment was carried out on the focal species *A. calendula*, *G. berardiana*, *H. glutinosum*, *H. glabra*, *P. aroides*, *P. debilis*, *P. canescens*, *T. cyanopetala*, *T. ornata*, *V. rosea* and *W. acuminata* in 2018 and 2019. Pairs of free-draining germination trays were deployed across a gradient of canopy cover in mid-October of both years, filled with soil, which had either been collected from the field and heat-sterilised to render pre-existing seeds nonviable (2019), or simply collected from the roadside (2018). Each germination tray consisted of 24 cells, with two cells randomly assigned to each focal species. In each cell, 15 seeds of the designated focal species were broadly distributed and lightly misted with water to facilitate seed-soil contact and minimise removal by wind. Trays placed in 2018 used seeds collected at the end of the 2017 growing season and dry after-ripened at 60°C for a month before being stored in cool, dry conditions at the University of Queensland. Seeds planted in 2019 were collected at the end of the 2018 growing season and were placed directly from the field into the germination trays. To re-establish microbial communities for those trays where the soil had been heat-treated, seeds were lightly covered with a small amount of untreated soil collected from the site in which they were buried. Untreated soil was collected from directly underneath coarse woody debris in patches where it was present as prior research in this system has shown that the effect of coarse woody debris on plant performance is partially attributable to debris-specific soil microbial communities (A. Pastore unpublished data). Some of the trays received the additions of leaf litter, but the results of this treatment were not included for the seed rates used in this study. Seed germination rate was measured by counting the number of seedlings which emerged in the field, but seed survival rate was not calculated in this experiment.

## 7 Supplementary Results

**Figure 1:**
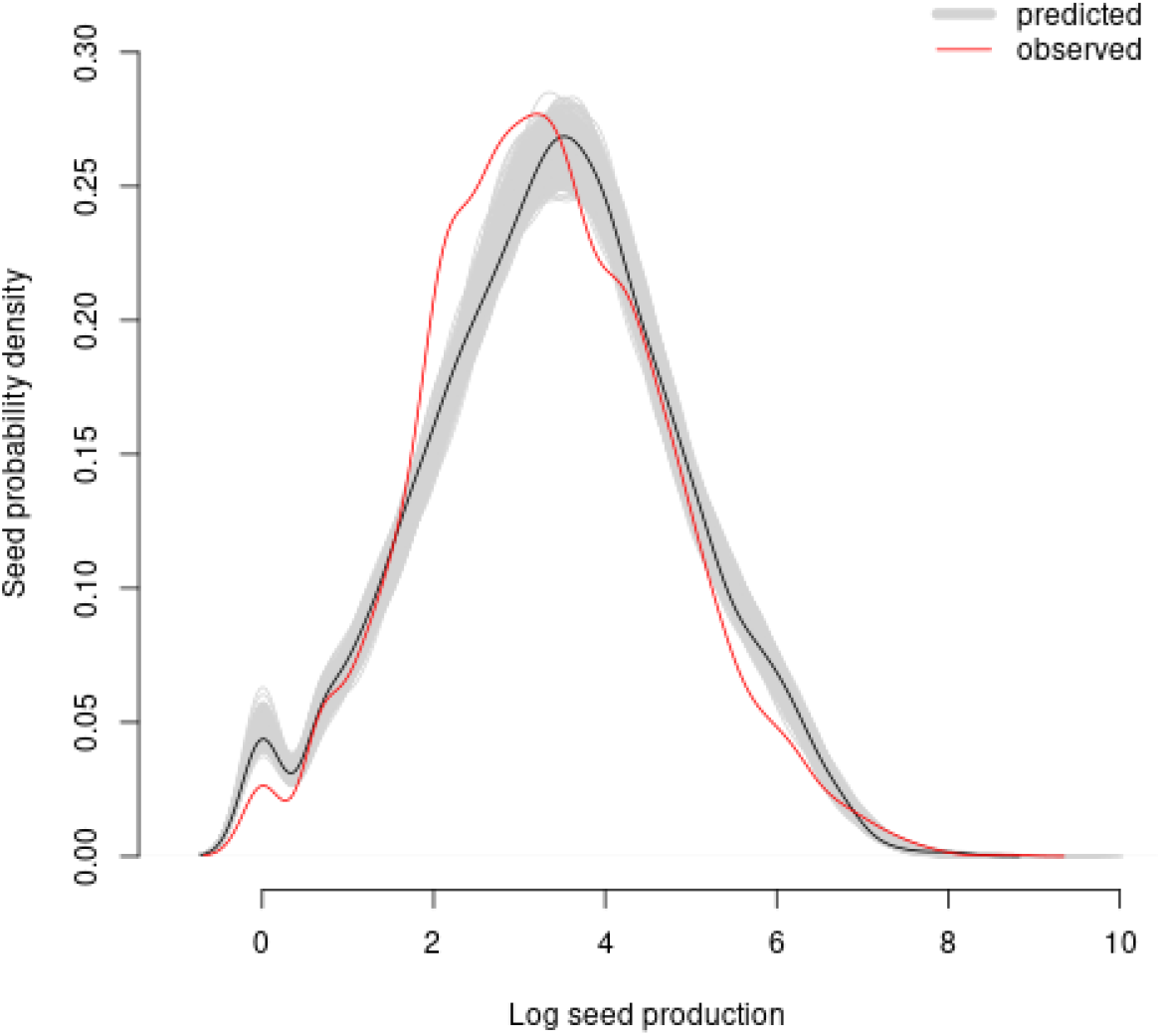
Posterior predictive check showing the density distribution of observed seed production values (red line) to simulated seed production values (light grey) as estimated by the RIM only, on a log scale. Simulated values were generated using the 80% posterior confidence intervals for each parameter, the black line shows simulated values using the median of each parameter.

**Figure 2:**
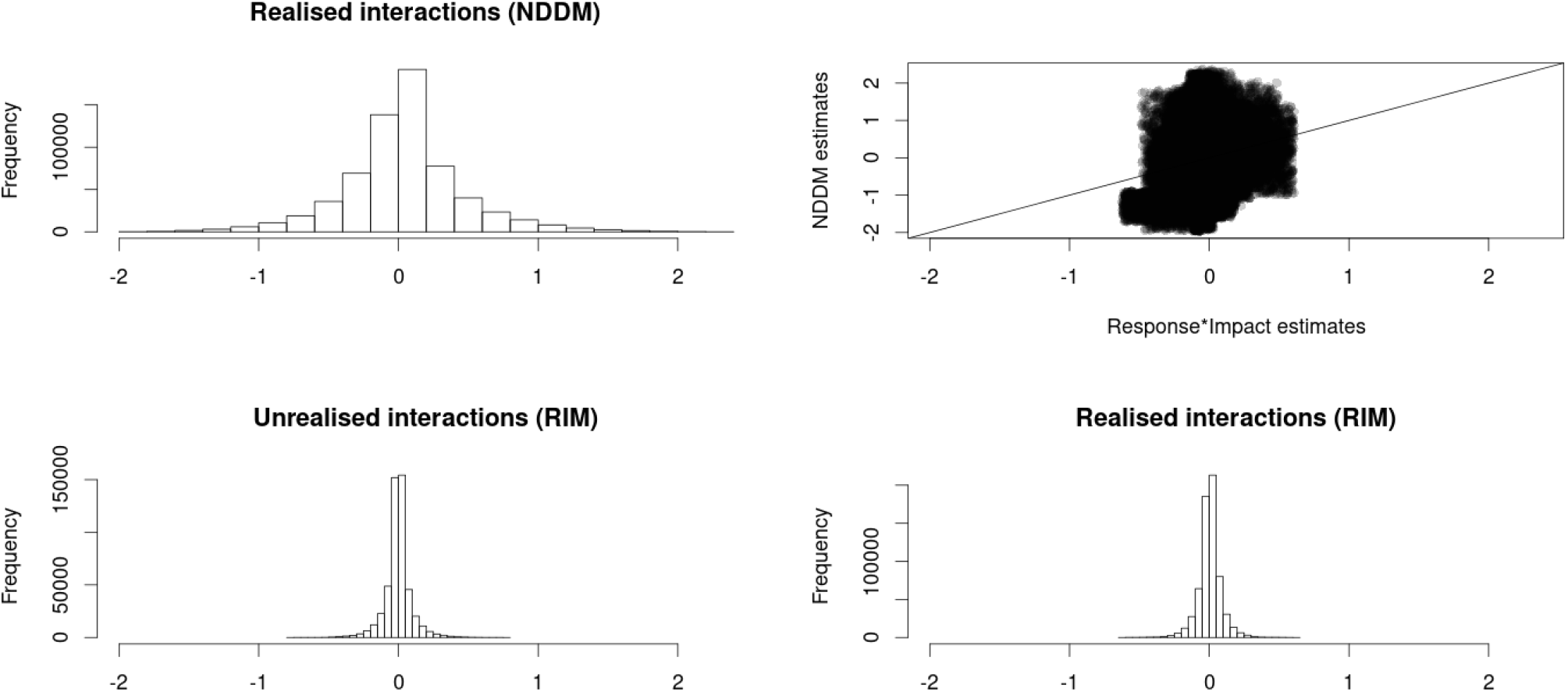
Distribution of interaction estimates from our case study. Parameter estimates are sampled from the 80% posterior confidence intervals returned by STAN. Upper left panel shows the distribution of observed interactions as estimated by the NDDM (*β_d_k_,j_*), which are then plotted against the corresponding RIM estimates (*r_d_k__e_j_*, x-axis) in the upper right panel. Bottom rows show the distribution of *unidentifiable* (left) and identifiable interactions estimates returned by the RIM. Interaction estimates are unscaled.

**Figure 3:**
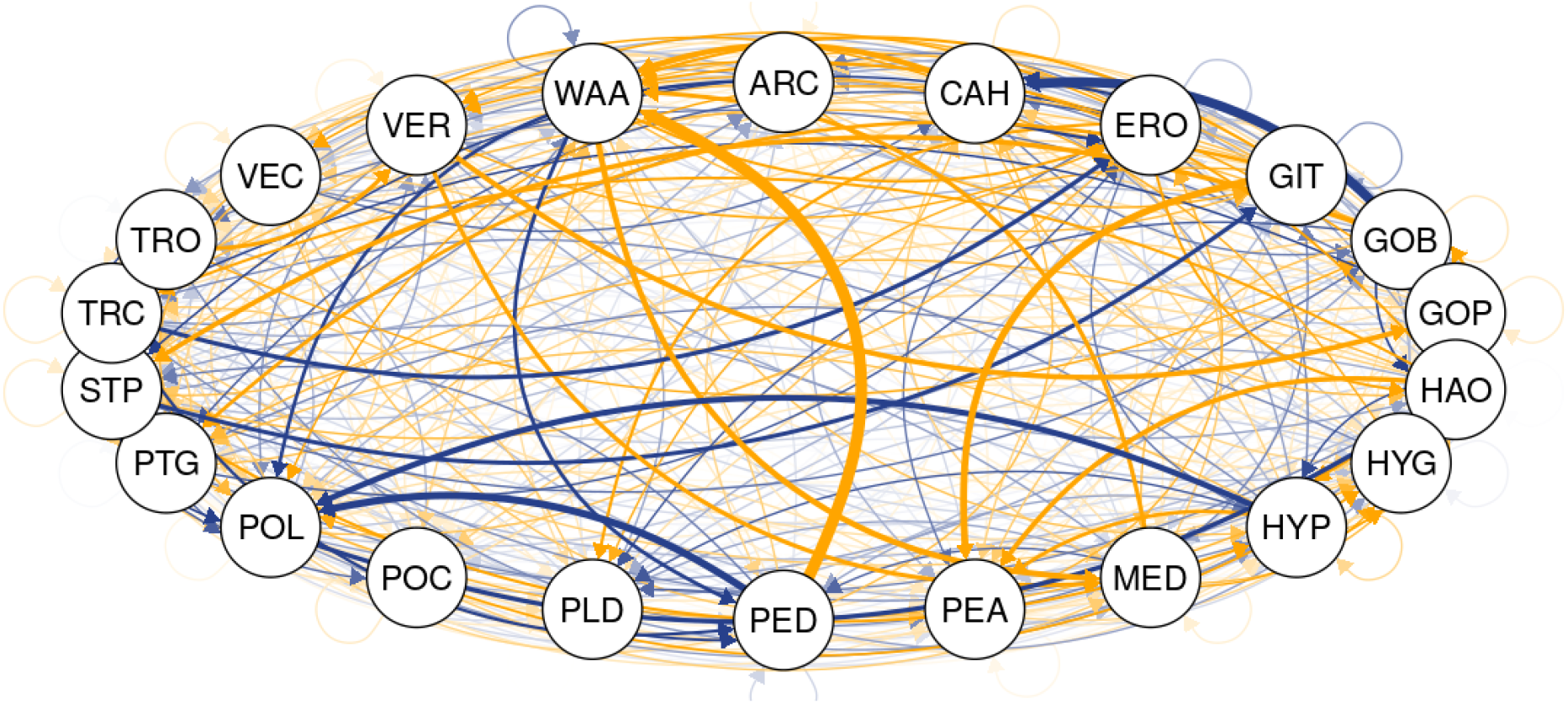
The full network of unscaled interactions between 22 focal species as estimated by the joint modelling framework presented here. Arrows point to species *i* and line thickness denotes median interaction strength over 1000 samples drawn from the 80% posterior confidence interval of each parameter. Facilitative interactions are shown in blue and competitive interactions are shown in yellow.

**Figure 4:**
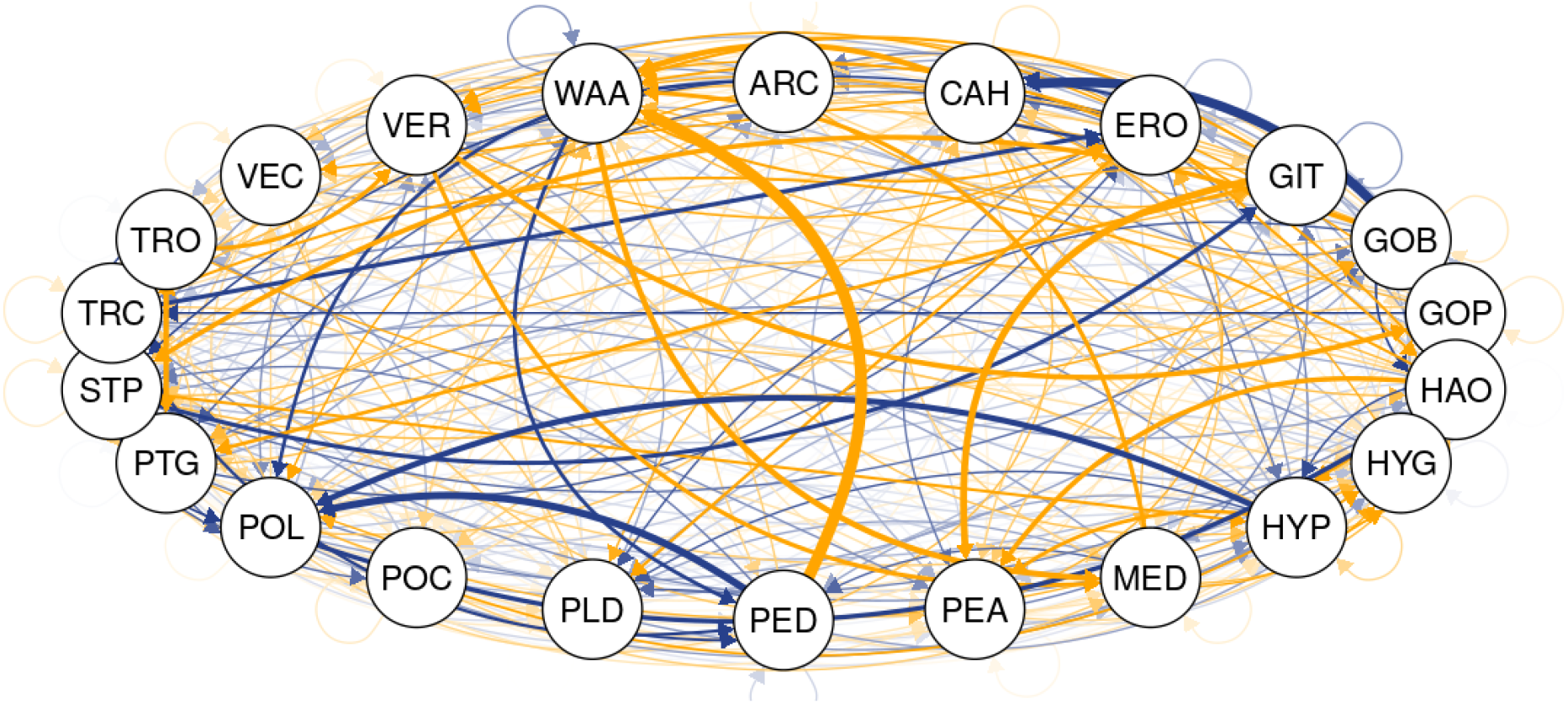
Network of identifiable interactions (unscaled) between 22 focal species as estimated by the NDDM, unidentifiable interactions are not shown here. Arrows point to species *i* and line thickness denotes median interaction strength over 1000 samples drawn from the 80% posterior confidence interval of each parameter. Facilitative interactions are shown in blue and competitive interactions are shown in yellow.

**Figure 5:**
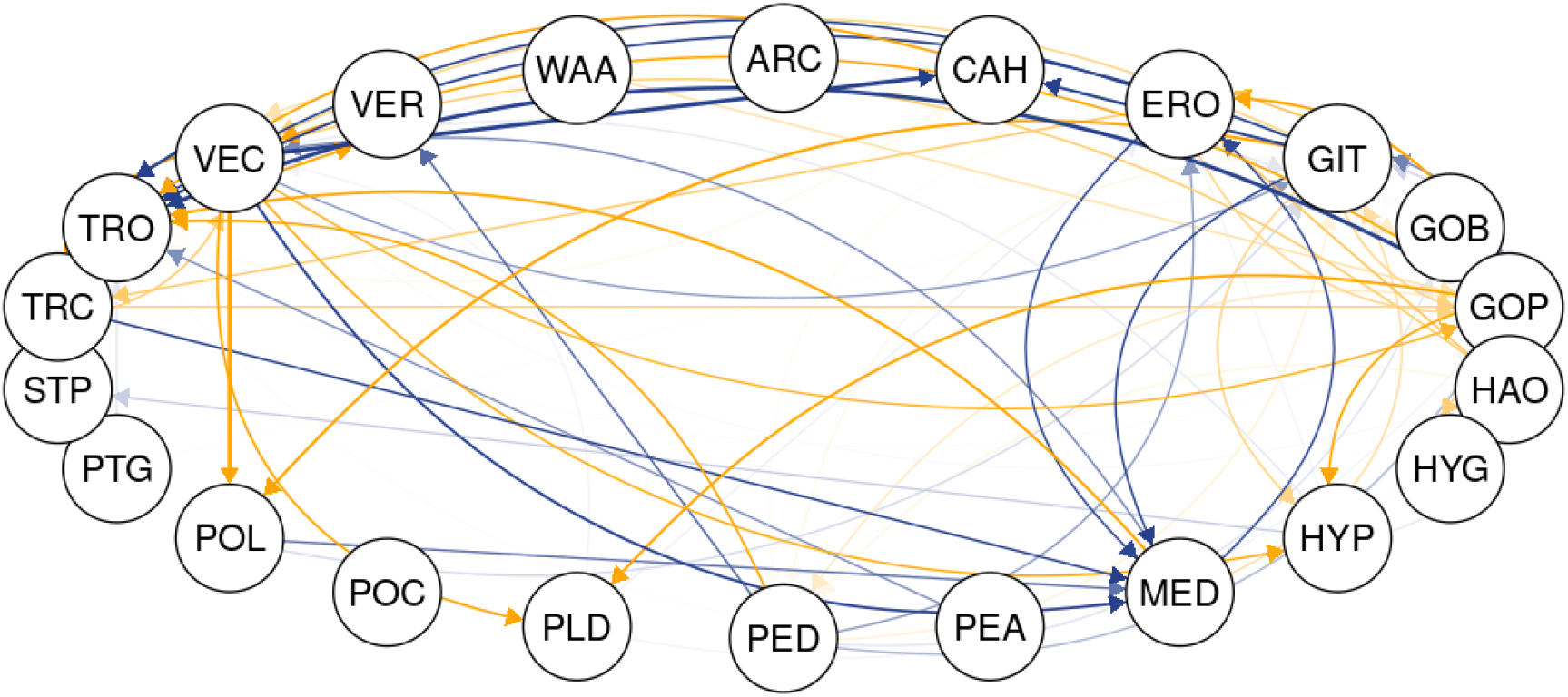
Network of unidentifiable interactions (unscaled) between 22 focal species as estimated by the RIM. Arrows point to species *i* and line thickness denotes median interaction strength over 1000 samples drawn from the 80% posterior confidence interval of each parameter. Facilitative interactions are shown in blue and competitive interactions are shown in yellow.

## Notes

https://github.com/malbion/JointModelFramework

### Competing Interest Statement

The authors have declared no competing interest.

## References

Barabás, G., Michalska-Smith, M. J., and Allesina, S. (2017). “Self-regulation and the stability of large ecological networks”. In: Nature Ecology & Evolution 1.12, pp. 1870–1875. DOI: 10.1038/s41559-017-0357-6.

Barner, A. K. et al. (2018). “Fundamental contradictions among observational and experimental estimates of non-trophic species interactions”. In: Ecology 0.0, pp. 1–10. DOI: 10.1002/ecy.2133.

Bascompte, J. and Melian, C. J. (2005). “Simple trophic modules for complex food webs”. In: Ecology 86.11, pp. 2868–2873. DOI: 10.1890/05-0101.

Bersier, L.-F., Banasek-Richter, C., and Cattin, M.-F. (2002). “Quantitative descriptors of food-web matrices”. In: Ecology 83.9, pp. 2101–2117.

Bimler, M. D. et al. (2018). “Accurate predictions of coexistence in natural systems require the inclusion of facilitative interactions and environmental dependency”. In: Journal of Ecology 106.5, pp. 1839–1852. DOI: 10.1111/1365-2745.13030.

Blanchet, F. G., Cazelles, K., and Gravel, D. (2020). “Co-occurrence is not evidence of ecological interactions”. In: Ecology Letters 23.7, pp. 1050–1063. DOI: 10.1111/ele.13525.

Broekman, M. J., Jongejans, E., and Tuljapurkar, S. (2020). “Relative contributions of fixed and dynamic heterogeneity to variation in lifetime reproductive success in kestrels (Falco tinnunculus)”. In: Population Ecology 62.4, pp. 408–424. DOI: 10.1002/1438-390X.12063.

Brooker, R. W. et al. (2008). “Facilitation in plant communities: The past, the present, and the future”. In: Journal of Ecology 96.1, pp. 18–34. DOI: 10.1111/j.1365-2745.2007.01295.x.

Bruno, J. F., Stachowicz, J. J., and Bertness, M. D. (2003). “Inclusion of facilitation into ecological theory”. In: Trends in Ecology and Evolution 18.3, pp. 119–125. DOI: 10.1016/S0169-5347(02)00045-9.

Burns, K. C. and Zotz, G. (2010). “A hierarchical framework for investigating epiphyte assemblages: networks, meta-communities, and scale”. In: Ecology 91.2, pp. 377–385.

Callaway, R. M. et al. (1997). “Competition and Facilitation: A Synthetic Approach to Interactions in Plant Communities”. In: Ecology 78.7, pp. 1958–1965. DOI: 10.1890/0012-9658(1997)078[1958:CAFASA]2.0.CO;2.

Carpenter, B. et al. (2017). “Stan: A probabilistic programming language”. In: Journal of Statistical Software 76.1. DOI: 10.18637/jss.v076.i01.

Chu, C. and Adler, P. B. (2015). “Large niche differences emerge at the recruitment stage to stabilize grassland coexistence”. In: Ecological Monographs 85.3, pp. 373–392. DOI: 10.1890/14-1741.1.

Cirtwill, A. R., Dalla Riva, G. V., et al. (2018). “A review of species role concepts in food webs”. In: Food Webs 16, e00186. DOI: 10.1016/j.fooweb.2018.e00093.

Cirtwill, A. R. and Stouffer, D. B. (2015). “Knowledge of predator–prey interactions improves predictions of immigration and extinction in island biogeography”. In: Global Ecology and Biogeography 25.7, pp. 900–911. DOI: 10.1111/geb.12332.

Connell, J. H. (1961). “The influence of interspecific competi-tion and other factors on the distribution of the barnacle Chthamalus stellatus”. In: Ecology 42, pp. 710–723.

Corbin, J. D. and D’Antonio, C. M. (2004). “Competition between native perenial and exotic annual grasses:implications for an historical invasion”. In: Ecology 85.5, pp. 1273–1283. DOI: 10.1890/02-0744.

Coyte, K. Z., Schluter, J., and Foster, K. R. (2015). “The ecology of the microbiome: Networks, competition, and stability”. In: Science 350.6261, pp. 663–666. DOI: 10.1126/science.aad2602.

Delmas, E. et al. (2019). “Analysing ecological networks of species interactions”. In: Biological Reviews 94.1, pp. 16–36. DOI: 10.1111/brv.12433.

Dunne, J. a., Lafferty, K. D., et al. (Jan. 2013). “Parasites affect food web structure primarily through increased diversity and complexity.” In: PLoS biology 11.6, e1001579. DOI: 10.1371/journal.pbio.1001579.

Dunne, J. a., Williams, R. J., and Martinez, N. D. (2002). “Network structure and biodiversity loss in food webs: Robustness increases with connectance”. In: Ecology Letters 5.4, pp. 558–567. DOI: 10.1046/j.1461-0248.2002.00354.x.

Ellison, A. M. (1996). “An introduction to Bayesian inference for ecological research and environmental decision-making”. In: Ecological Applications 6.4, pp. 1036–1046.

Ellison, A. M. (2004). “Bayesian inference in ecology”. In: Ecology Letters 7.6, pp. 509–520. DOI: 10.1111/j.1461-0248.2004.00603.x.

Ellison, A. M. (2019). “Foundation Species, Non-trophic Interactions, and the Value of Being Common”. In: iScience 13, pp. 254–268. DOI: 10.1016/j.isci.2019.02.020.

Fisher, R. A., Corbet, A. S., and Williams, C. B. (1943). “The Relation Between the Number of Species and the Number of Individuals in a Random Sample of an Animal Population”. In: The Journal of Animal Ecology 12.1, p. 42. DOI: 10.2307/1411.

Geweke, J. (1992). “Evaluating the accuracy of sampling-based approaches to calculating posterior moments”. In: Bayesian Statistics 4. Ed. by J. Bernado et al. Oxford: Clarendon Press, pp. 169–193.

Giling, D. P. et al. (2019). “Plant diversity alters the representation of motifs in food webs”. In: Nature Communications 10.1, pp. 1–7. DOI: 10.1038/s41467-019-08856-0.

Godoy, O., Bartomeus, I., et al. (2018). “Towards the Integration of Niche and Network Theories”. In: Trends in Ecology & Evolution 33.4, pp. 287–300. DOI: 10.1016/j.tree.2018.01.007.

Godoy, O., Kraft, N. J. B., and Levine, J. M. (2014). “Phylogenetic relatedness and the determinants of competitive outcomes”. In: Ecology Letters 17.7, pp. 836–844. DOI: 10.1111/ele.12289.

Godoy, O. and Levine, J. M. (2014). “Phenology effects on invasion success: Insights from coupling field experiments to coexistence theory”. In: Ecology 95.3, pp. 726–736. DOI: 10.1890/13-1157.1.

Gorman, E. J. O. et al. (2019). “A simple model predicts how warming simplifies wild food webs”. In: Nature Climate Change 9, pp. 611–616. DOI: 10.1038/s41558-019-0513-x.

Grace, J. B. and Tilman, D. (1990). Perspectives on plant competition. Academic Press, p. 484.

Griffith, D. M., Veech, J. A., and Marsh, C. J. (2016). “cooccur: Probabilistic Species Co-Occurrence Analysis in R”. In: Journal of Statistical Software 69, pp. 1–17. DOI: 10.18637/jss.v069.c02.

Gross, N. et al. (2015). “Functional equivalence, competitive hierarchy and facilitation determine species coexistence in highly invaded grasslands”. In: New Phytologist 206.1, pp. 175–186. DOI: 10.1111/nph.13168.

Hammill, E. et al. (2015). “Food web persistence is enhanced by non-trophic interactions”. In: Oecologia 178.2, pp. 549–556. DOI: 10.1007/s00442-015-3244-3.

Huber, P., Ronchetti, E., and Victoria-Feser, M.-P. (2004). “Estimation of Generalized Linear Latent Variable Models”. In: Journal of the Royal Statistical Society. Series B (Statistical Methodology) 66.4, pp. 893–908.

Hui, F. K. C. (2021). boral: Bayesian Ordination and Regression AnaLysis.

Jordano, P. (2016). “Sampling networks of ecological interactions”. In: Functional Ecology 30.12, pp. 1883–1893. DOI: 10.1111/1365-2435.12763.

Kidziński, Ł. et al. (2021). “Generalized Matrix Factorization”. In: arXiv preprint.

Kinlock, N. L. (2019). “A Meta-Analysis of Plant Interaction Networks Reveals Competitive Hierarchies as Well as Facilitation and Intransitivity”. In: American Naturalist 194.5. DOI: 10.1086/705293.

Lafferty, K. D. et al. (2008). “Parasites in food webs: The ultimate missing links”. In: Ecology Letters 11.6, pp. 533–546. DOI: 10.1111/j.1461-0248.2008.01174.x.

Laird, R. A. and Schamp, B. S. (2006). “Competitive Intransitivity Promotes Species Coexistence”. In: The American Naturalist 168.2, pp. 182–193. DOI: 10.1086/506259.

Laska, M. S. and Wootton, J. T. (1998). “Theoretical concepts and empirical approaches to measuring interaction strength”. In: Ecology 79.2, pp. 461–476. DOI: 10.1890/0012-9658(1998)079[0461:TCAEAT]2.0.CO;2.

Levine, J. M. and HilleRisLambers, J. (2009). “The importance of niches for the maintenance of species diversity.” In: Nature 461.7261, pp. 254–257. DOI: 10.1038/nature08251.

Libralato, S., Christensen, V., and Pauly, D. (2006). “A method for identifying keystone species in food web models”. In: Ecological Modelling 195.3-4, pp. 153–171. DOI: 10.1016/j.ecolmodel.2005.11.029.

Losapio, G., Cruz, M. de la, et al. (2018). “The assembly of a plant network in alpine vegetation”. In: Journal of Vegetation Science 29.6, pp. 999–1006. DOI: 10.1111/jvs.12681.

Losapio, G. and Schöb, C. (2017). “Resistance of plant–plant networks to biodiversity loss and secondary extinctions following simulated environmental changes”. In: Functional Ecology 31.5, pp. 1145–1152. DOI: 10.1111/1365-2435.12839.

Martin, A. D., Quinn, K. M., and Park, J. H. (2011). “MCMCpack: Markov Chain Monte Carlo in R”. In: Journal of Statistical Software 42.9, p. 22.

Martyn, T. E. (2020). “Understanding the role of direct and indirect interactions in mediating local plant diversity”. PhD Thesis. University of Queensland, p. 246. DOI: https://doi.org/10.14264/uql.2020.824.

Mayfield, M. M. and Stouffer, D. B. (2017). “Higher-order interactions capture unexplained complexity in diverse communities”. In: Nature Ecology & Evolution 1, pp. 1–7. DOI: 10.1038/s41559-016-0062.

Miele, V. et al. (2019). “Non-trophic interactions strengthen the diversity — functioning relationship in an ecological bioenergetic network model”. In: PLoS Computational Biology, pp. 1–20.

Montesinos-Navarro, A. et al. (2018). “Community structure informs species geographic distributions”. In: PLoS ONE 13.5, pp. 1–16. DOI: 10.1371/journal.pone.0197877.

Naeem, S. et al. (2000). “Plant diversity increases resistance to invasion in the absence of covarying extrinsic factors”. In: Oikos 91.1, pp. 97–108.

Narwani, A. et al. (2019). “Interactive effects of foundation species on ecosystem functioning and stability in response to disturbance”. In: Proceedings of the Royal Society B 286.

Niku, J. et al. (2021). “Analyzing environmental-trait interactions in ecological communities with fourth-corner latent variable models”. In: Environmetrics 32.6, e2683. DOI: 10.1002/env.2683.

Novak, M. and Wootton, J. T. (2010). “Using experimental indices to quantify the strength of species interactions”. In: Oikos 119.7, pp. 1057–1063. DOI: 10.1111/j.1600-0706.2009.18147.x.

Nyakatya, M. J. and McGeoch, M. A. (2008). “Temperature variation across Marion Island associated with a keystone plant species (Azorella selago Hook. (Apiaceae))”. In: Polar Biology 31.2, pp. 139–151. DOI: 10.1007/s00300-007-0341-8.

Olesen, J. M. et al. (2011). “Missing and forbidden links in mutualistic networks”. In: Proceedings of the Royal Society B: Biological Sciences 278.1706, pp. 725–732. DOI: 10.1098/rspb.2010.1371.

Orwin, K. H. et al. (2016). “Season and dominant species effects on plant trait-ecosystem function relationships in intensively grazed grassland”. In: Journal of Applied Ecology 55, pp. 236–245. DOI: 10.1111/1365-2664.12939.

Paine, R. T. (1969). “A Note on Trophic Complexity and Community Stability”. In: The American Naturalist 103.929, pp. 91–93. DOI: 10.1086/282586.

Paine, R. T. (1992). “Food-web analysis through field measurement of per capita interaction strength”. In: Nature 355.6355, pp. 73–75. DOI: 10.1038/355073a0.

Picoche, C. and Barraquand, F. (2020). “Strong self-regulation and widespread facilitative interactions in phytoplankton communities”. In: Journal of Ecology 108.6, pp. 2232–2242. DOI: 10.1111/1365-2745.13410.

Pimm, S. L. and Lawton, J. H. (1978). “On feeding on more than one trophic level”. In: Nature 275.5680, pp. 542–544. DOI: 10.1038/275542a0.

Piraino, S. et al. (2002). “Variability of species’ roles in marine communities : change of paradigms for conservation priorities”. In: Marine Biology 140.5, pp. 1067–1074. DOI: 10.1007/s00227-001-0769-2.

Plummer, M. et al. (2006). “CODA: Convergence Diagnosis and Output Analysis for MCMC”. In: R News 6.1, pp. 7–11.

Power, M. E. et al. (1996). “Challenges in the Quest for Keystones”. In: BioScience 46.8, pp. 609–620. DOI: 10.2307/1312990.

R Core Team (2020). R: A Language and Environment for Statistical Computing. Vienna, Austria.

Ramus, A. P. et al. (2017). “An invasive foundation species enhances multifunctionality in a coastal ecosystem”. In: PNAS 114.32, pp. 1–6. DOI: 10.1073/pnas.1700353114.

Riley, L. A., Dybdahl, M. F., and Hall, R. O. (2008). “Invasive species impact: Asymmetric interactions between invasive and endemic freshwater snails”. In: Journal of North American Benthological Society 27.3, pp. 454–520. DOI: 10.1899/07.

Rodriguez, L. F. (2006). “Can invasive species facilitate native species? Evidence of how, when, and why these impacts occur”. In: Biological Invasions 8.4, pp. 927–939. DOI: 10.1007/s10530-005-5103-3.

Ross, W. Y. F. C. et al. (2011). “Ecosystem ecology meets adaptive management : food web response to a controlled flood on the Colorado River, Glen Canyon”. In: Ecological Applications 21.6, pp. 2016–2033.

Saiz, H. et al. (2018). “The structure of plant spatial association networks is linked to plant diversity in global drylands”. In: Journal of Ecology 106.4, pp. 1443–1453. DOI: 10.1111/1365-2745.12935.

Sander, E. L., Wootton, J. T., and Allesina, S. (2017). “Ecological Network Inference from Long-Term Presence-Absence Data”. In: Scientific Reports 7.1, pp. 1–12. DOI: 10.1038/s41598-017-07009-x.

Soulé, M. E. et al. (2005). “Strongly interacting species: conservation policy, management, and ethics”. In: BioScience 55.2, pp. 168–176.

Stan Development Team (2020). RStan: the R interface to Stan. R package version 2.19.3.

Stouffer, D. B., Cirtwill, A. R., and Bascompte, J. (Sept. 2014). “How exotic plants integrate into pollination networks”. In: Journal of Ecology 102. Ed. by I. Bartomeus, pp. 1442–1450. DOI: 10.1111/1365-2745.12310.

Thompson, R. M. et al. (Dec. 2012). “Food webs: reconciling the structure and function of biodiversity.” English. In: Trends in ecology & evolution 27.12, pp. 689–97. DOI: 10.1016/j.tree.2012.08.005.

Thurman, L. L. et al. (2019). “Testing the link between species interactions and co-occurrence in a trophic network”. In: Ecography 42, pp. 1658–1670. DOI: 10.1111/ecog.04360.

Ulanowicz, R. E. and Wolff, W. F. (1991). “Ecosystem flow networks: Loaded dice?” In: Mathematical Biosciences 103.1, pp. 45–68. DOI: 10.1016/0025-5564(91)90090-6.

Wainwright, C. E., Holt, R. D., and Mayfield, M. M. (2019). “Looks can be deceiving: ecologically similar exotics have different impacts on a native competitor”. In: Oecologia 190.4, pp. 927–940. DOI: 10.1007/s00442-019-04468-z.

Wootton, J. T. and Emmerson, M. (2005). “Measurement of Interaction Strength in Nature”. In: Annual Review of Ecology, Evolution, and Systematics 36.1, pp. 419–444. DOI: 10.1146/annurev.ecolsys.36.091704.175535.

Yenni, G., Adler, P. B., and Ernest, S. K. M. (2017). “Do persistent rare species experience stronger negative frequency dependence than common species?” In: Global Ecology and Biogeography 26, pp. 513–523. DOI: 10.1111/geb.12566.

Yenni, G., Adler, P. B., and Ernest, S. K. M. (2012). “Strong self-limitation promotes the persistence of rare species”. In: Ecology 93.3, pp. 456–461.

Zhao, Q. et al. (2019). “Horizontal and vertical diversity jointly shape food web stability against small and large perturbations”. In: Ecology Letters 22.7, pp. 1152–1162. DOI: 10.1111/ele.13282.

Zheng, Y. et al. (2015). “Are invasive plants more competitive than native conspecifics? Patterns vary with competitors”. In: Scientific Reports 5.15622. DOI: 10.1038/srep15622.

